# Environmental stress sensitivity determines bacterial Mn(II) oxidation

**DOI:** 10.1101/2025.05.09.653071

**Authors:** Yaohui Bai, Hui lin, Qiaojuan Wang, Yuhan Wang, Jinsong Liang, Xue Ning, Jiuhui Qu

## Abstract

Bacterial Mn(II) oxidation is a widespread and ecologically significant process, with Mn oxides playing essential roles in biogeochemical cycling and environmental remediation. However, the environmental cues that initiate this microbial activity remain enigmatic. In this study, a spontaneous mutant was isolated from the previously characterized *Arthrobacter* strain that exhibited autonomous Mn(II)-oxidizing capacity, in contrast to the parental strain, which required cocultivation with other bacteria to initiate Mn(II) oxidation^1^. Comparative transcriptomic analysis suggested that a gene system encoding an environmental stress-sensing receptor controlled the transcription of *boxA*, a Mn(II)-oxidizing gene. These findings indicate that two conditions are required for bacterial Mn(II) oxidation: the presence of a Mn(II)-oxidizing gene and an environmentally responsive regulatory system capable of activating its expression. Based on this knowledge, eight additional bacterial strains were isolated that triggered *Arthrobacter* Mn(II) oxidation. Transcriptomic profiling revealed a robust positive correlation between *boxA* expression and genes encoding two-component signal transduction systems. Given their widespread distribution (approximately 23% of environmental bacterial genomes harbor Mn(II)-oxidizing gene homologs^2^) and constant exposure to diverse biotic and abiotic stressors, these bacteria are likely to undergo frequent Mn(II) oxidation triggered by microbial interactions or chemical signals. This study provides critical mechanistic insight into the environmental regulation of bacterial Mn(II) oxidation, addressing a longstanding gap in understanding its natural occurrence.

## Introduction

Manganese (Mn), the second most abundant transition metal in the Earth’s crust, plays a pivotal role in regulating biological activity and environmental redox processes. In natural ecosystems, soluble Mn(II) is readily oxidized into insoluble Mn(III/IV) oxide by a taxonomically diverse set of microorganisms, including both fungi and bacteria^3-7^. Fungal Mn(II) oxidation predominantly results in the formation of Mn(III)-organic complexes, in which trivalent manganese is stabilized through coordination with organic ligands. These complexes are critical for lignin depolymerization, enabling the degradation of this otherwise structurally recalcitrant organic polymer^8,9^. In contrast, bacterial Mn(II) oxidation preferentially generates Mn(IV) oxides, the thermodynamically favored oxidation state under aerobic conditions^10^. These Mn(IV) oxides are ubiquitous in the environment, accumulating in metal-contaminated streams, hydrothermal vent deposits, and across nearly one-third of the Pacific Ocean seafloor^11-14^. Beyond their geochemical prevalence, these oxides are hypothesized to serve multiple ecological and physiological roles, functioning as electron acceptors in anaerobic respiration, contributing to microbial energy conservation, and offering protection against ultraviolet radiation, oxidative damage, and biological predation^15,16^.

Interestingly, bacterial Mn(II) oxidation is often restricted to the stationary phase of growth^15^. Moreover, during our experiments, it revealed that some Mn(II)-oxidizing bacteria, such as *Pseudomonas*, display variable Mn(II) oxidation over time. These experimental results suggest that its induction is conditional. However, the precise environmental or physiological triggers that activate Mn(II)-oxidizing genes remain unresolved. Current evidence indicates that bacterial Mn(II) oxidation can occur via direct enzymic reactions^17,18^ or via an indirect process mediated by extracellular superoxide radical (O_2_^−^•), which non-enzymatically oxidizes Mn(II) ^19-21^ (this process is controversial). Identified Mn(II)-oxidizing enzymes belong to two principal protein families: animal heme peroxidases (AHPs) and multicopper oxidases (MCOs)^17,22-25^. These enzymes typically catalyze the oxidation of Mn(II) to Mn(IV) via a transient Mn(III) intermediate^17,26,27^. AHPs involved in Mn(II) oxidation have been characterized in multiple alphaproteobacterial species^17-19^ and in the gammaproteobacterium *Pseudomonas putida* GB-1^28^, while MCOs have been implicated in Mn(II) oxidation across a wide range of taxa, including *Leptothrix, Pseudomonas, Pedomicrobium*, and several marine spore-forming *Bacillus* species^17,22,23^. In addition to individual bacteria, microbe-microbe interactions have emerged as potent drivers of Mn(II) oxidation^1,29-31^. Notably, a recently uncovered syntrophic partnership between *Candidatus Manganitrophus noduliformans* and *Ramlibacter lithotrophicus* revealed a tightly coordinated system linking Mn(II) oxidation with autotrophic CO_2_ fixation^32^. Moreover, an electrosyntrophic interaction between *Rhodopseudomonas palustris* and *Geobacter metallireducens* has been shown to catalyze Mn(II) oxidation under anoxic, light-deprived conditions^33^. Collectively, these findings suggest that interspecies interactions may constitute a widespread and ecologically significant mechanism driving Mn(II) oxidation across diverse natural settings.

Despite the widespread presence of Mn oxides in natural environments and their critical roles in biogeochemical cycling and contaminant remediation, the biological relevance and regulatory mechanisms underlying bacterial Mn(II) oxidation remain poorly defined. In our previous study^1^, we speculated that Mn(II) oxidation in *Arthrobacter* sp. QXT-31 (hereafter *Arthrobacter*) was triggered by stress induced through co-culture with *Sphingopyxis* sp. QXT-31 (hereafter *Sphingopyxis*). To investigate this hypothesis and evaluate its broader relevance, a spontaneous *Arthrobacter* mutant capable of autonomous Mn(II) oxidation was used as a reference strain (Supplementary Fig. S1). Results demonstrated that environmental stress sensitivity, rather than the presence of Mn(II)-oxidizing gene *boxA* alone, constrained the initiation of Mn(II) oxidation. In addition, we found Mn(II) oxidation was frequently activated through interspecies interactions involving bacteria that harbored Mn(II)-oxidizing genes. These findings clarify the conditional nature of bacterial Mn(II) oxidation and highlight its ecological significance in microbial communities and Mn cycling.

## Results

### Environmental stress sensitivity regulates bacterial Mn(II) oxidation

Our previous study^1^ have reported *Sphingopyxis* triggered the expression of Mn(II)-oxidizing gene (*boxA*) of *Arthrobacter* via a stress-relevant interaction. To reveal the role of environmental stress on triggering enzymatic Mn(II) oxidation, co-culture experiments were conducted by combining *Arthrobacter* with varying concentrations of *Sphingopyxis*, using cell number ratios ranging from 4:1 to 1:4□096 to create a biological stress gradient for *Arthrobacter*. Results (Fig. 1a) showed Mn(II) oxidation was detected only when the ratio of *Sphingopyxis* to *Arthrobacter* exceeded 1:4. At higher *Sphingopyxis* proportions, such as 1:1□024 and 1:4□096, the onset of Mn(II) oxidation was significantly delayed. An optimum ratio between the two bacterial strains was identified that minimized the time required for Mn(II) oxide formation. These results imply that low stress levels are insufficient to induce Mn(II) oxidation, while excessive stress delays or suppresses the process. These observations support the hypothesis that Mn(II) oxidation by *Arthrobacter* is a stress-driven process and that the timing of its activation depends on stress intensity.

**Figure 1.**
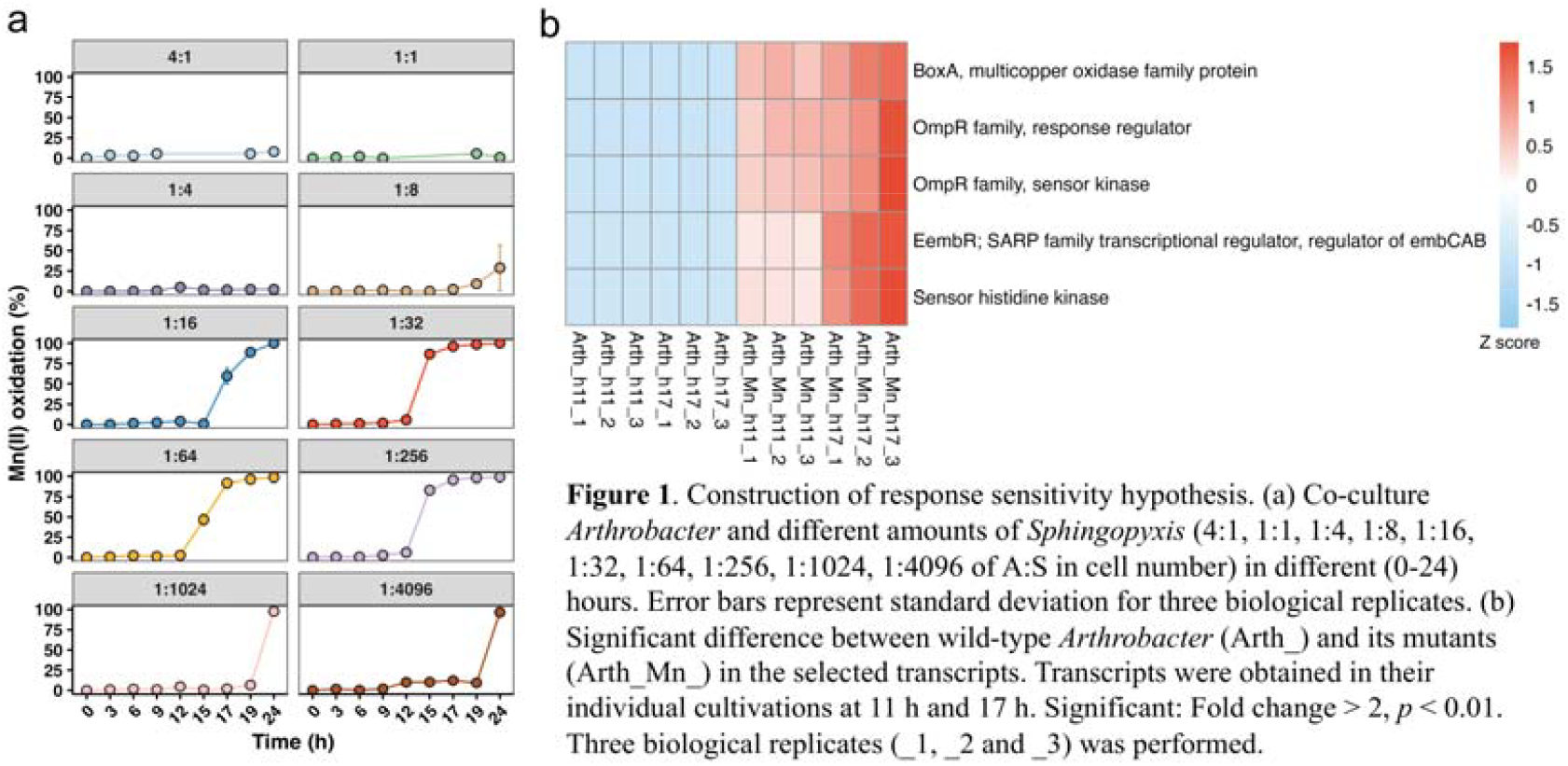
Construction of response sensitivity hypothesis. (a) Co-culture Arthtobacter and different amounts of Sphingopyxis (4:1, 1:1, 1:4, I 1:16, 1:32, 1:64, 1:256, l: 1024, 1:4096 of A:S in cell number) in different (0-24) hours. Error bars represent standard deviation for three biological replicates. (b) Significant difference between wild-type Arthrobacter (Arth_) and its mutants (Arth Mn_) in the selected transcripts. Transcripts were obtained in their individual cultivations at 11 h and 17 h. Significant: Fold change > 2, *p* < 0.01. Three biological replicates 2 and _3) was performed.

To verify this hypothesis, a spontaneous *Arthrobacter* mutant capable of Mn(II) oxidation in monoculture was isolated and purified (Fig. S1). Transcriptomic comparisons between the mutant and wild-type strains across different time points (Fig. S2 and Supplementary Excel file: Transcript comparison) identified genes co-upregulated with the Mn(II)-oxidizing gene *boxA*. Both *boxA* and two genes encoding a two-component system (TCS) showed significant upregulation at the onset of Mn(II) oxidation (Fig. 1b). In contrast, these genes were not obviously expressed in the wild-type strain, suggesting that TCS activation is required to initiate *boxA* transcription. As TCS modules mediate adaptive responses to diverse extracellular and intracellular stimuli^3^□, the finding of TCS genes upregulation in the mutant demonstrate that variation in stress response sensitivity within genetically similar backgrounds governs the initiation of bacterial Mn(II) oxidation.

### Mechanistic link between environmental stress sensitivity and bacterial Mn(II) oxidation

In the Mn(II)-oxidizing *Arthrobacter* mutant, transcript abundance of the TCS genes was substantially higher than that of the Mn(II)-oxidizing gene *boxA* (Fig. 1b). As such, we thought that TCS genes and *boxA* would have a greater separation, with higher levels of TCS transcripts to achieve effective activation of *boxA* expression. Genomic analysis revealed that both the wild-type and mutant strains possess genomes of approximately 5 Mbp. Based on gene position data from the wild-type *Arthrobacter* genome (Supplementary Excel file: Transcript comparison, start and stop sites), the two upregulated TCS genes were found to be adjacent, with one encoded on the forward strand and the other on the reverse strand. As anticipated, these TCS genes (start site 347□210) were located at a distant from the Mn(II)-oxidizing gene *boxA* (start site 516□350) (Fig. 2a). Comparative genomic analysis between *Arthrobacter* and its Mn(II)-oxidizing mutant revealed no sequence variation in these loci (Supplementary Excel File: Genome SNP comparison), nor detectable differences in DNA methylation (Supplementary Excel file: Methylation comparison). These results indicate that the enhanced sensitivity to environmental stress observed in the mutant is not attributable to genetic mutations or epigenetic modifications in the relevant genes, although the underlying mechanism remains unresolved.

**Figure 2.**
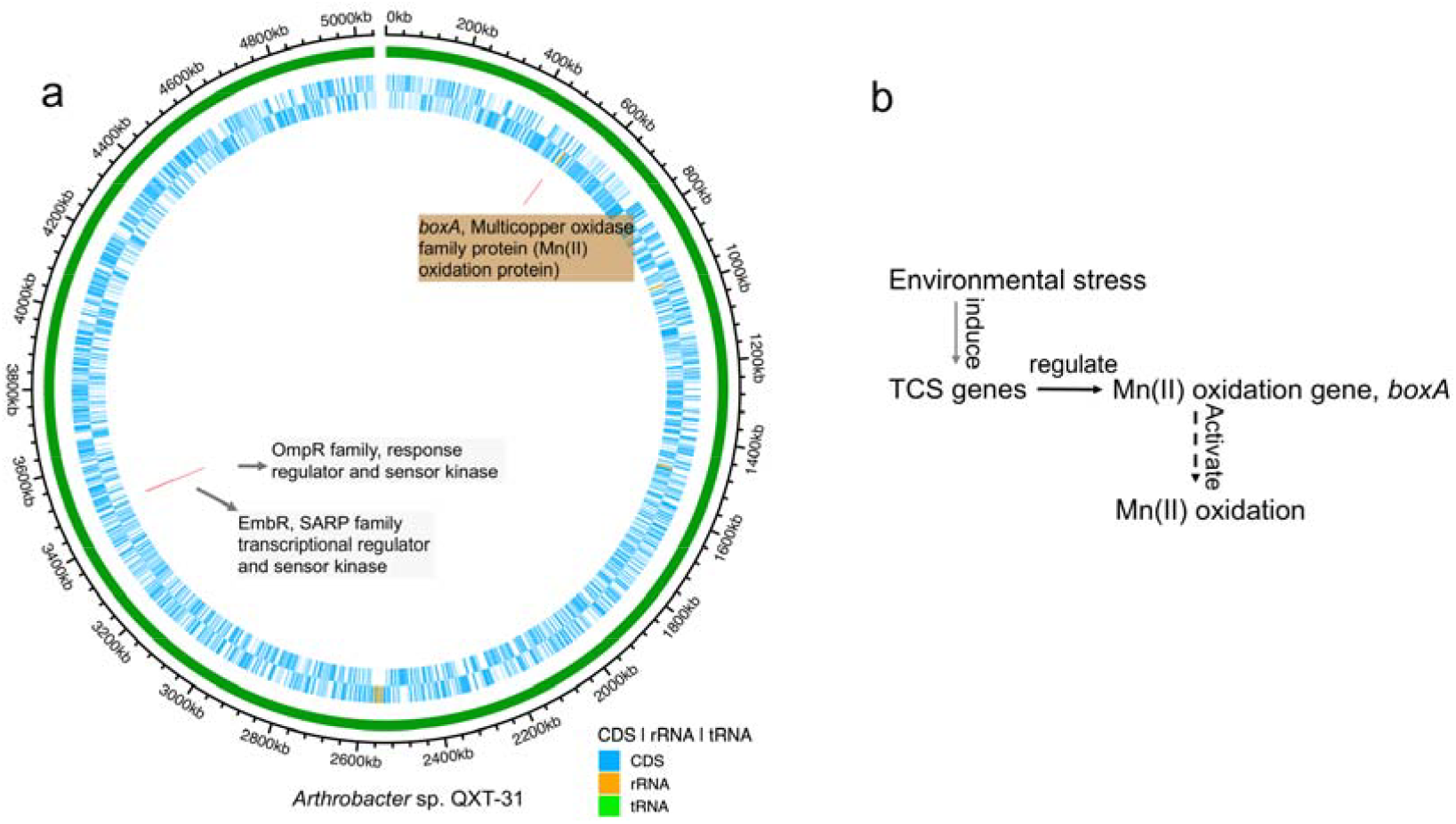
Environmental stress sensitivity determines Mn(II) oxidation by *Arthrobacter*. (a) Genomic comparison between *Arthrobacter* and its Mn(II)-oxidizing mutant; (b) Proposed regulatory mechanism of Mn(II) oxidation by *Arthrobacter*.

Based on these findings, a regulatory model for stress-induced bacterial Mn(II) oxidation was proposed (Fig. 2b). Upon exposure to environmental stressors such as nutrient deprivation (inspired by the observation that bacterial Mn(II) oxidation is often restricted to the stationary phase of growth^15^) or antagonistic microbial factors, Mn(II)-oxidizing bacteria activated the TCS signaling cascade. When TCS transcript levels surpass a threshold, downstream regulatory pathways are triggered, leading to the induction of *boxA* and other stress-responsive genes, including those encoding antioxidant enzymes that maintain redox homeostasis. The coordinated transcriptional activation of these pathways culminates in the onset of Mn(II) oxidation.

### Environmental stress sensitivity determines occurrence of bacterial Mn(II) oxidation in natural settings

To evaluate whether environmental stress-induced Mn(II) oxidation represents a broader ecological phenomenon, bacterial isolates were obtained from soil and screened for their ability to induce Mn(II) oxidation in *Arthrobacter* during co-culture. Eight distinct bacterial strains were identified that consistently triggered Mn(II) oxidation when paired with *Arthrobacter* (Table S1). Phylogenetic analysis revealed that these strains, along with the previously studied *Sphingopyxis*, were taxonomically affiliated with Alpha- and Gammaproteobacteria (Fig. 3a). Transcriptomic analysis showed a positive correlation between *boxA* expression and transcripts encoding components of the TCS (Fig. 3b and Fig. S4a), suggesting that these partner strains imposed environmental stress sufficient to activate the Mn(II)-oxidizing response in *Arthrobacter*. Notably, peroxidase gene expression also increased with *boxA* expression (Fig. S4b), consistent with a broader stress response aimed at mitigating oxidative damage. Additional stress-inducing agents were also tested, including eight bacterial consortia (Table S2 and Fig. S5), a single-celled eukaryote (Fig. S6), and two antibiotic compounds (Fig. S7). In each case, Mn(II) oxidation in *Arthrobacter* was successfully induced, providing strong support for the hypothesis that a wide range of natural stressors can trigger Mn(II) oxidation by *Arthrobacter*.

**Figure 3.**
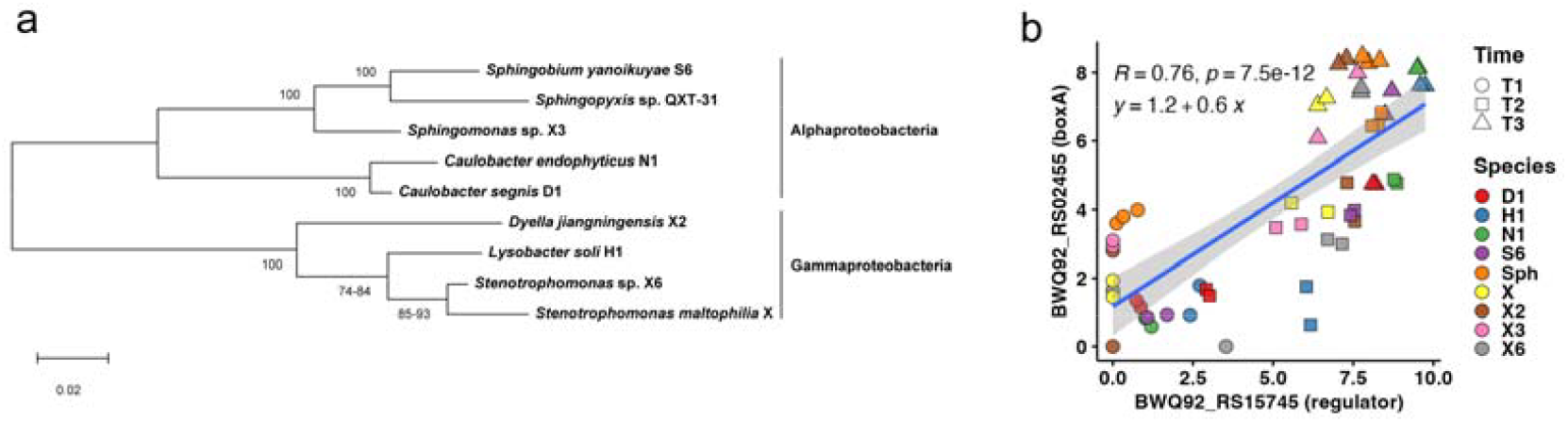
Induction of Mn(II) oxidation in *Arthrobacter* by co-cultured bacterial strains. (a) Phylogenetic affiliations of eight bacterial isolates that activated Mn(II) oxidation. (b) Correlation between *Arthrobacter boxA* transcripts and the TCS regulator (embCAB) across three sampling points (T1, T2, and T3). Details are described in Supplementary Excel file: Correlation of transcripts.

## Discussion

Bacterial Mn(II) oxidation plays a pivotal role in geochemical cycling and environmental engineering, yet its biological significance and regulatory mechanisms remain poorly understood. This gap limits both the predictive understanding of Mn biotransformation and the development of effective environmental remediation strategies. To address this, a combination of culture-based assays and transcriptomic analyses was employed to investigate the triggering conditions for Mn(II) oxidation. Results revealed that environmental stress was sensed through the bacterial TCS, which subsequently activated the transcription of Mn(II)-oxidizing genes. These findings establish TCS expression as a key regulatory node in bacterial Mn(II) oxidation, consistent with prior observations in *P. putida* GB-1^34^. However, the earlier study did not clarify the involvement of MCOs, which are widely recognized as key catalytic components in this process. The expression of TCS appears to be governed by two factors: the magnitude of external environmental stress and the intrinsic sensitivity of the bacterial strain to such stimuli. This dual dependency suggests two general modes of Mn(II) oxidation. In the first, a single Mn(II)-oxidizing strain must both encode a functional oxidase gene (e.g., MCO) and exhibit sufficient sensitivity to environmental stressors such as nutrient limitation. In the second, interspecies interactions act as the trigger: one strain possesses the genetic capacity for Mn(II) oxidation, while a second strain(s) imposes stress—such as the production of virulence factors in the case of *Sphingopyxis*^1^—sufficient to induce the oxidative response in the Mn(II)-oxidizing partner. This mechanism may explain the frequent activation of Mn(II) oxidation in natural microbial communities, where taxonomically diverse strains co-exist and interact through a broad range of biochemical signals and stressors.

In summary, this study demonstrated that bacterial Mn(II) oxidation is a stress-responsive process shaped by both genetic potential and environmental context and is frequently initiated through interactions with other microbial or chemical stressors in natural settings. These findings also raise a deeper question: is Mn(II) oxidation—or the accumulation of Mn(IV) oxides—a strategy to preserve the sensitivity and homeostatic balance of stress-responsive bacteria? Resolving this question will pave the way to find the physiological significance of Mn(II) oxidation in microbial life.

## Materials and Methods

### Bacterial strains and media

*Arthrobacter* sp. QXT-31 and *Sphingopyxis* sp. QXT-31 strains used in this study were described in a previous study ^1^. The *Arthrobacter* strain harbors a Mn(II)-oxidizing gene but exhibits no Mn(II) oxidation under standard conditions, whereas the *Sphingopyxis* strain induces Mn(II) oxidation in *Arthrobacter* during co-culture. Monocultures of both strains were initially cultured in PYG medium^1^ (10 g of tryptone, 10 g of NaCl, 2 g of glucose, and 5 g of yeast extract per liter) in the absence of Mn(II).

### Mn(II) oxidation under varying *Arthrobacter* and *Sphingopyxis* ratios

To assess the effect of cell ratio on Mn(II) oxidation, *Arthrobacter* and *Sphingopyxis* strains were co-inoculated into fresh medium at various ratios (A:S of 4:1, 1:1, 1:4, 1:8, 1:16, 1:32, 1:64, 1:256, 1:1□024, 1:4□096) and incubated at 30°C with shaking at 170 rpm overnight. Cell counts during co-cultivation were quantified using viable plate counts. Cultures were homogenized by vigorous shaking, serially diluted (10^5^– 10^9^-fold), and immediately plated on solid PYG medium. Colonies were differentiated by morphology: white for *Arthrobacter* and yellow for *Sphingopyxis*. Colony-forming units (CFUs) were determined to assess cell densities for both strains. All experiments were performed with three biological replicates.

### Isolation and identification of Mn(II)-oxidizing *Arthrobacter* mutant

The *Arthrobacter* strain was cultivated on solid PYG medium containing 5.5 mg/L Mn(II). After approximately one month, weak Mn(II) oxidation was observed. Colonies showing oxidation were transferred to liquid PYG medium containing 5.5 mg/L Mn(II), incubated for 48 h, and re-plated on solid PYG medium containing 5.5 mg/L Mn(II). This procedure was repeated twice. A mutant capable of autonomous Mn(II) oxidation was successfully isolated and designated *Arthrobacter* sp. QXT-31 mutant (CGMCC No. 32889, Fig. S1). Cell biomass was monitored by measuring optical density at 600□nm (OD_600_).

### RNA extraction, sequencing, and transcriptomic analysis

Monocultures of each bacterial strain were cultivated for 24 hours in the presence of 100 μM MnCl□, followed by mixing for co-cultivation. Samples were collected at 0, 3, and 6 hours after the onset of co-culture. Cell pellets were harvested by centrifugation at 10□000 *g* for 5 min at 4°C and immediately subjected to RNA extraction using TRNzol Reagent (TIANGEN, Beijing, China) according to the manufacturer’s instructions. RNA integrity was evaluated using an Agilent 2100 Bioanalyzer (Agilent Technologies, Santa Clara, CA, USA), and only samples with RNA integrity number (RIN) or RNA quality number (RQN) values greater than 8.0 were selected for cDNA library construction and sequencing. Libraries were sequenced on the Illumina HiSeq 2500 platform, generating 150-bp paired-end reads.

Sequencing reads were mapped to the reference genome sequences using Bowtie2^35^. Mapped reads were then counted using featureCounts^36^ and subsequently analyzed for differential expression using the DEseq2^37^ algorithm in R (v4.4.1). For pathway enrichment analysis, KEGG Orthology (KO) identifiers were assigned to all reference genes using KofamScan ^38^ with the KOfam-HMM profiles database (v2024-08-01). KO identifiers from both the reference genome and differentially expressed genes (DEGs) were used for enrichment analysis with the R package clusterProfiler (v4.0.2) ^39^.

### Comparative genomic analysis of *Arthrobacter* and its mutant

Whole-population genome sequencing was conducted to identify genetic alterations between *Arthrobacter* and its Mn(II)-oxidizing mutant. Sequencing libraries were generated using a NEBNext® Ultra™ DNA Library Prep Kit for Illumina (NEB, USA) following the manufacturer’s recommendations. Index barcodes were added to assign reads to each sample. Sequencing was performed on the Illumina NovaSeq 6000 platform, generating 150-bp paired-end reads. Multiple quality control parameters, including read quality and GC content, were assessed using FastQC (v0.11). Low-quality reads (Phred score < 20) were trimmed using Cutadapt (v2.10) ^40^. High-quality reads were then aligned against the reference genome using the Burrows-Wheeler Aligner (BWA, v0.7.4)^41^ under standard parameters. Alignment files were then sorted and indexed using SAMtools (v1.3.1). Single nucleotide polymorphisms (SNPs) and insertions/deletions (InDels) were identified using Breseq (v0.30.0)^42^, while structural variants (SVs) were detected using BreakDancer and VarScan (v2.4.0)^43^. The annotated reference genome for *Arthrobacter* QXT-31 (GenBank: CP019304.1) was retrieved from NCBI and used to annotate the identified polymorphisms.

For methylome profiling, DNA from the mutant strain was submitted to Beijing Novogene Bioinformatics Technology Co., Ltd. (China) for single-molecule real-time (SMRT) sequencing (PacBio Core Enterprise). SMRTbell DNA template libraries were prepared according to the Pacific Biosciences protocol for 10-kb template preparation and sequencing. This involved DNA fragmentation, DNA damage and end repair, blunt-end ligation, purification with 0.45× AMPure PB Beads, size selection via the BluePippin System, and post-selection damage repair. Library quality was assessed using a Qubit® 2.0 Fluorometer (Thermo Scientific) and fragment size was estimated using an Agilent 2100 Bioanalyzer (Agilent Technologies). The SMRTbell libraries were sequenced with a 120-min movie acquisition time using a P4 polymerase-C2 DNA sequencing reagent kit following standard PacBio RS II instrument protocols (Pacific Biosciences). Genome-wide analyses for base modification and motif detection were performed using the RS Modification and Motif Analysis protocol in SMRT Analysis v2.3.0 Patch 5. The FASTA reference genome sequence (*A*. QXT-31; CP019304.1) used for base modification detection was obtained from the NCBI database. A Circos plot was generated using the “circlize” package in R.

### Isolation of bacterial strains capable of inducing Mn(II) oxidation in

#### *Arthrobacter* during co-cultivation

Soil samples were collected from diverse environments, including grassland, forest, mountain, rhizosphere, and riverside habitats (Table S1), to isolate bacterial strains capable of inducing Mn(II) oxidation in *Arthrobacter*. Each soil sample was suspended in sterile water at a 1:4 (w/w) ratio, and a 24-hour pre-culture of *Arthrobacter* was added at a final volume ratio of 10% (v/v). The suspension was serially diluted and plated onto solid PYG medium supplemented with 100 μM Mn(II), followed by incubation at 30°C. Colonies capable of producing Mn oxides in spatial association with *Arthrobacter*, identified based on distinct colony morphology, were isolated by streak plating. Colony identity was confirmed via full-length 16S rRNA gene sequencing, aligned against the reference *Arthrobacter* 16S rRNA gene^1^ with criteria of 100% identity and 100% coverage. Mn(II) oxidation was further validated for each candidate strain in both solid and liquid PYG media supplemented with Mn(II).

For co-culture experiments, *Arthrobacter* and each isolated strain capable of inducing Mn(II) oxidation were grown independently for 24 hours, then mixed at a 1:1 volumetric ratio. The resulting co-cultures were used for Mn(II) oxidation monitoring, growth measurement (OD_600_), and RNA-seq. At specific time points, 1 mL of culture was sampled, and residual Mn(II) concentration was quantified to assess Mn(II) oxidation kinetics.

Genomic DNA from the isolated bacterial strains was extracted and subjected to next-generation sequencing on the Illumina HiSeq 4000 platform with 250-bp paired-end reads and an insert size of 400 bp, yielding approximately 2 Gbp of reads. Quality filtering was conducted using fastp (v0.23.1)^44^ with default parameters. Draft genomes were assembled using metaSPAdes (v1.15)^45^, and genome completeness and contamination were assessed using CheckM2^46^. Full-length 16S rRNA genes were predicted from the draft genome using Barrnap (https://github.com/tseemann/barrnap) and queried using BLASTN against the NCBI 16S rRNA sequence database. Taxonomic assignments were made at the species level for hits exhibiting ≥99% sequence identity.

### Phylogenetic tree construction of eight isolated strains

Full-length 16S rRNA gene sequences for the eight bacterial isolates (Table 1) were predicted from whole-genome assemblies using Barrnap (v0.9, https://github.com/tseemann/barrnap) with default parameters. Sequences were aligned using MUSCLE (v5.3, parameter: -super5) ^47^. The resulting alignments were trimmed using trimAl (v1.4) ^48^ and an unrooted phylogenetic tree was constructed using IQ-TREE (v2.0, parameters: -m MFP -T AUTO -bb 1000) ^49^. Maximum-likelihood phylogenetic analysis was performed in MEGA 12^50^. Bootstrap support values for each clade were derived from adaptive resampling and are shown at branch nodes.

### Determination of Mn(II) oxidation activity

Mn(II) oxidation was assessed by monitoring the decrease in soluble Mn(II) concentration. At each sampling point, cultures were cooled on ice and centrifuged at 10□000 *g* for 3 min at 4 °C. The resulting supernatant was carefully removed, taking care not to resuspend the settled solids, and filtered through a 0.45-μm membrane. Mn(II) concentrations in the filtrate were measured using ICP-OES (710, Agilent, USA).

## Data availability

All raw reads were deposited in the China National Center for Bioinformation database (https://www.cncb.ac.cn) under BioProject PRJCA038441.

## Acknowledgments

This work was supported by the National Natural Science Foundation of China (52450009 and 52388101). We also thank Peijun Zhang and Haiyang Li for their helps in the experiments.

## Notes

The authors declare no competing financial interest.

