## Supplemental Materials for "Environmental stress sensitivity determines bacterial Mn(II) oxidation"

**Figure S1. Growth and Mn(II) oxidation by the mutant of** ***Arthrobacter* sp. QXT-31 (hereafter *Arthrobacter*).** *Arthrobacter* sp. QXT-31-Mn (represents the mutant) colonies were aseptically transferred from a solid agar medium to Erlenmeyer flasks containing 50 mL of PYG medium without Mn²⁺ using a sterile inoculation loop. The culture was incubated at 30°C with shaking at 170 rpm for 12 hours until reaching an optical density at 600 nm (OD₆₀₀) of 0.8. Subsequently, 10% (v/v) of the activated culture was inoculated into fresh PYG medium supplemented with 100 µM Mn²⁺. Prior to inoculation, HEPES buffer was sterilized using a 0.22-μm membrane filter and added to the medium to a final concentration of 0.1 M. Culture aliquots were aseptically sampled at designated time intervals to measure soluble Mn²⁺ concentrations and OD₆₀₀. Data represent means of three biological replicates. Error bars denote standard deviation.

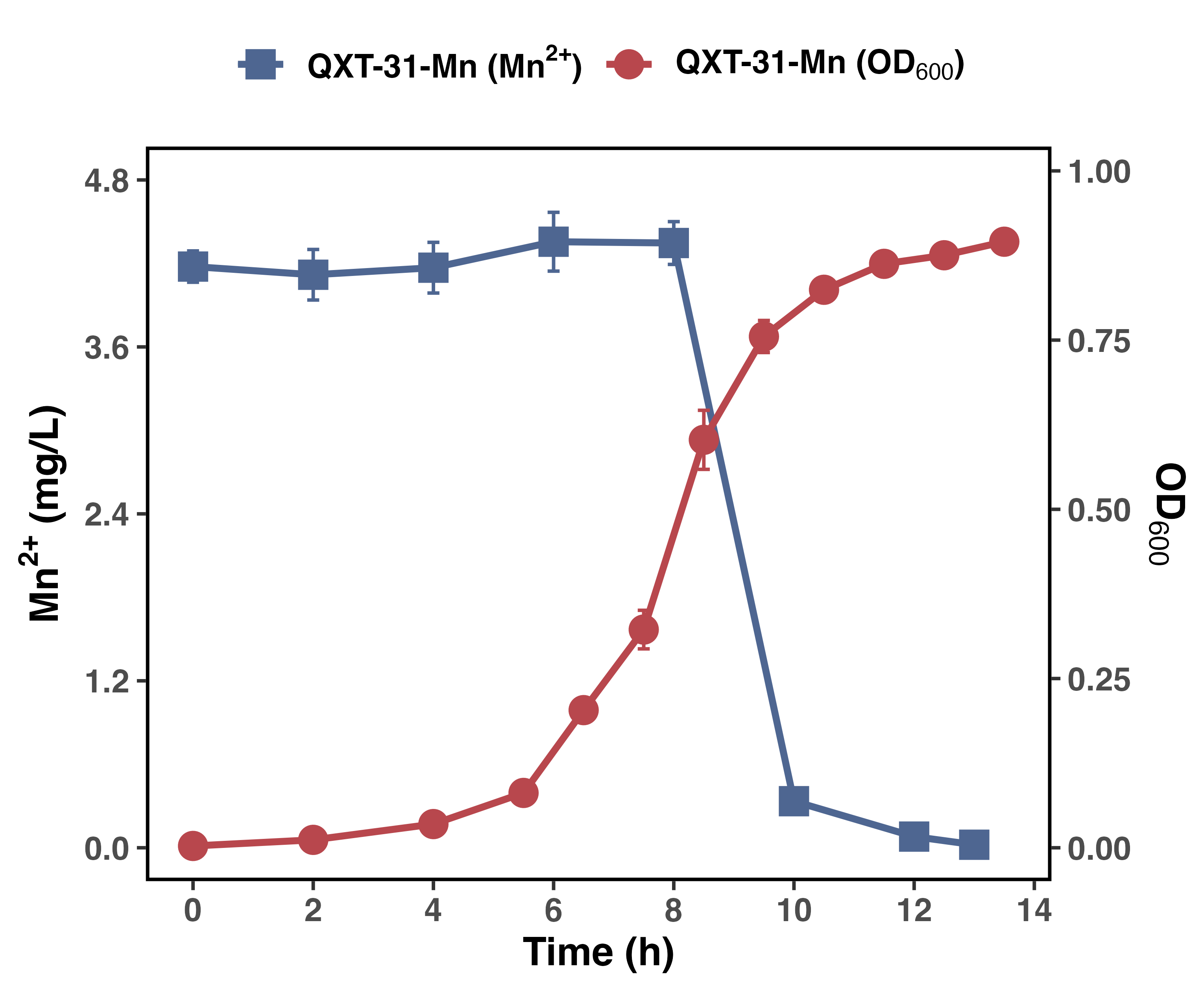

**Figure S2. Differential gene expression between wild-type *Arthrobacter* and its Mn(II)-oxidizing mutant.** Transcripts were obtained from monocultures of the wild-type and mutant strains at 11 and 17 hours. Differential expression analysis was performed using DESeq2 in R. Genes with fold-change > 2 and *p* < 0.01 were considered significantly differentially expressed.

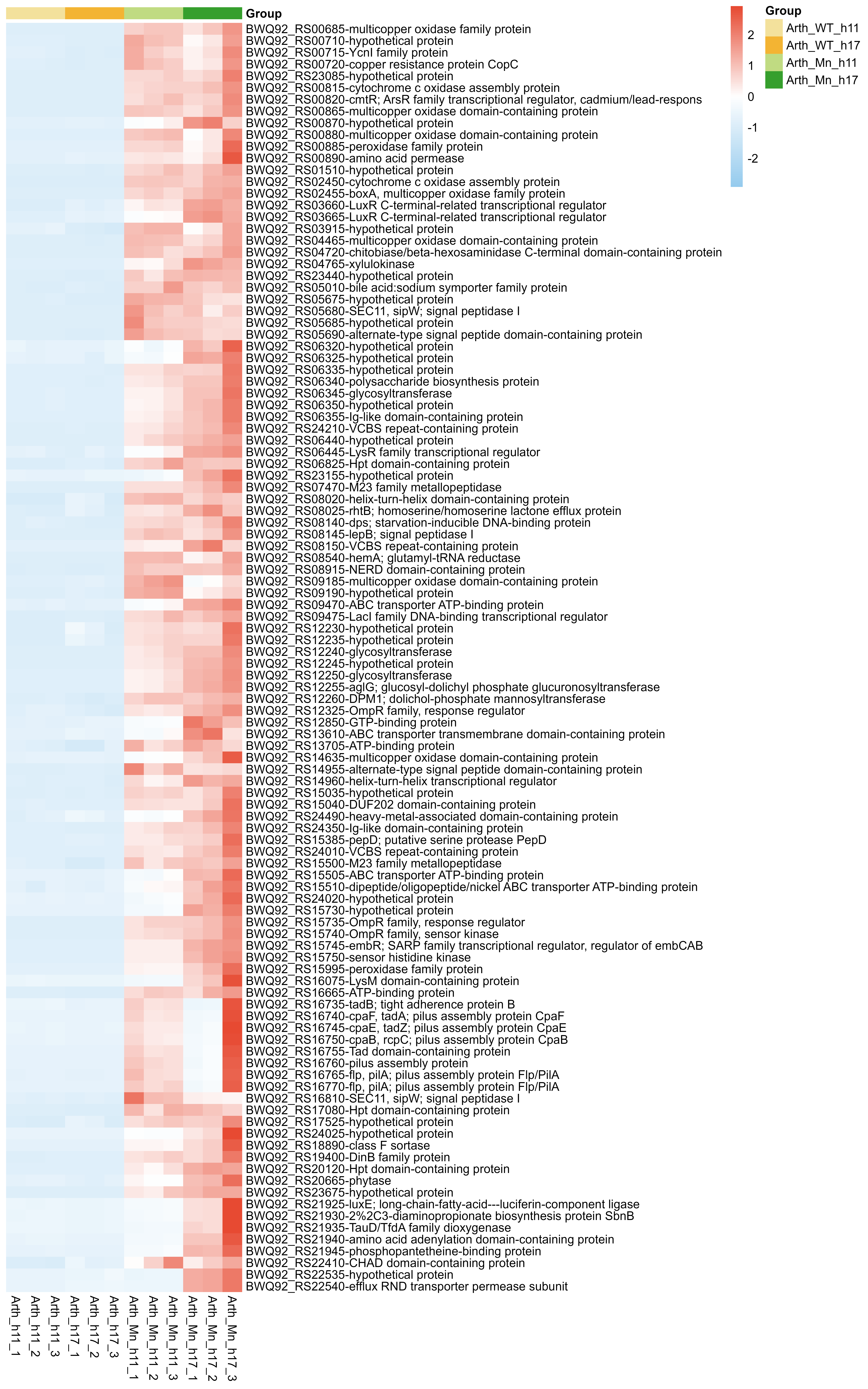

Z score

**Figure S3. Mn(II) oxidation and growth of *Arthrobacter* during co-culture with each of eight bacterial strains in solid and liquid PYG media.** For liquid medium experiments, data represent the means of three biological replicates; error bars indicate standard deviation.

**Solid medium**

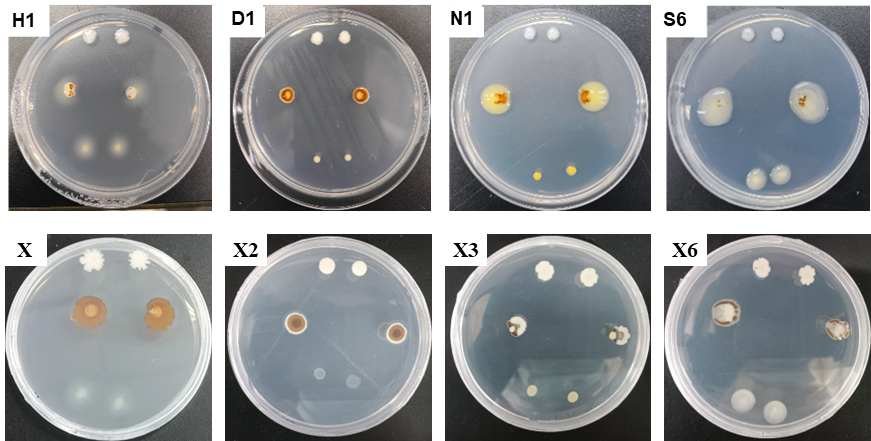

**Liquid medium**

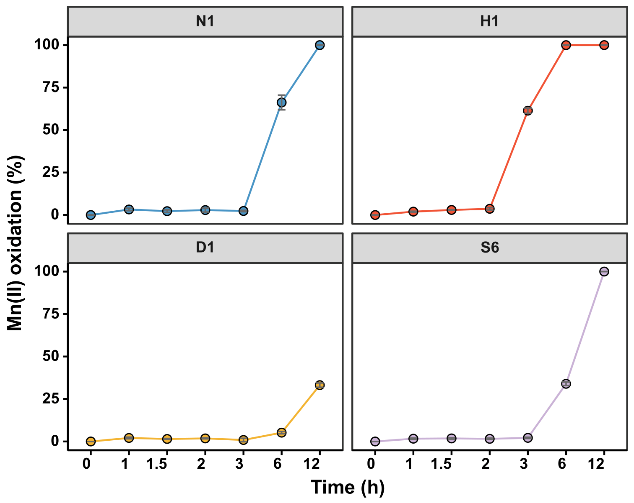

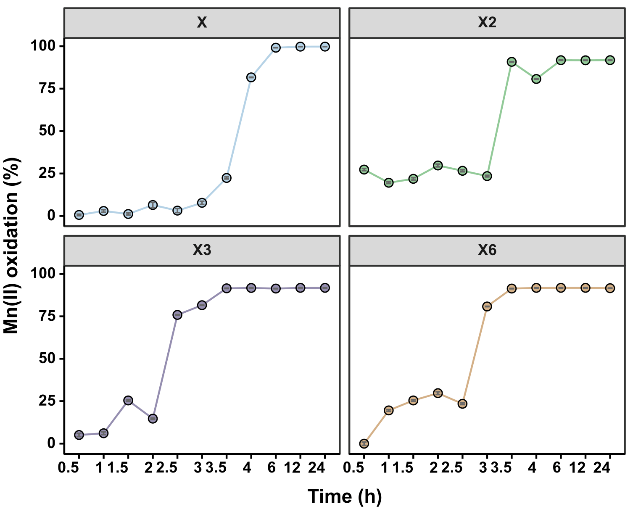

**Figure S4.** **Log-transformed transcript abundance correlations among two-component system (TCS), Mn(II) oxidation, and peroxidase genes.** (a) Correlation between transcripts encoding embCAB-associated TCS regulator and corresponding sensor histidine kinase. (b) Correlation between transcripts of Mn(II)-oxidizing *boxA* and peroxidase gene. To avoid distortions from zero values, transcript abundances were transformed using log₁₀(abundance + 1). T1, T2, and T3 represent three distinct sampling time points, which varied by strain. Additional details are provided in the Supplementary Excel file: Correlation of transcripts.

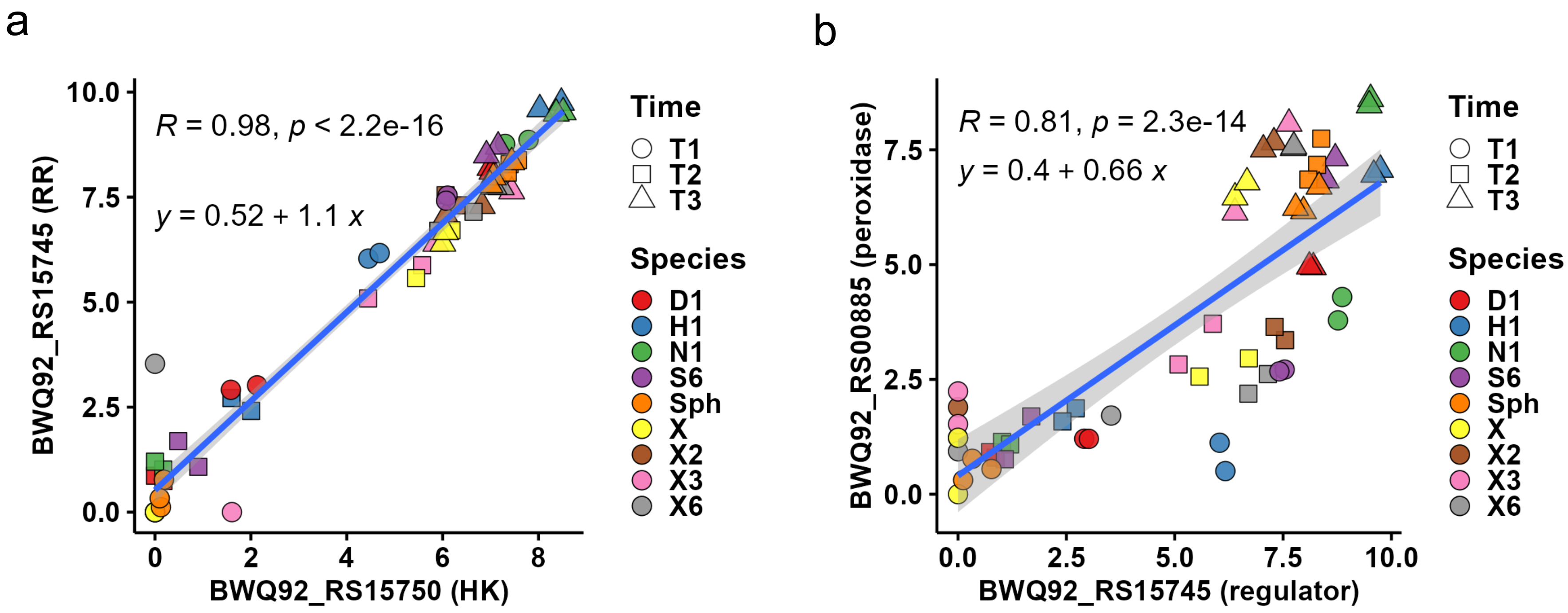

**Figure S5. Bacterial consortia that trigger Mn(II) oxidation in *Arthrobacter*.** Colonies capable of inducing Mn(II) oxidation in *Arthrobacter* were isolated from co-cultures with environmental samples. Full-length 16S rRNA gene sequencing revealed that these interacting partners were not single strains but bacterial consortia comprising multiple genera (listed in Table S2).

**Experimental procedure:** *Arthrobacter* sp. QXT-31 was cultured in PYG medium and co-inoculated with various soil samples. The mixtures were plated onto solid PYG medium containing 100 μM Mn(II) and incubated at 30°C. Mn(II) oxide formation was observed at the interaction zone between two distinct non-Mn(II)-oxidizing colonies. One colony, resembling *Arthrobacter* sp. QXT-31 in morphology, was confirmed as *Arthrobacter* via full-length 16S rRNA gene sequencing. Colonies adjacent to the Mn(II) oxide zone were isolated for purification through successive streaking. Following multiple purification cycles, putatively axenic cultures were processed for DNA extraction, and full-length 16S rRNA genes were amplified using universal primers 27F and 1492R ^1^. Sanger sequencing and BLASTN analysis against the NCBI 16S rRNA database confirmed one isolate as *Arthrobacter*, while persistent mixed chromatograms from the interacting partner indicated the presence of multiple bacterial species. To resolve this taxonomic complexity, metagenomic sequencing was performed, followed by genome assembly and contig binning to reconstruct draft genomes for each member of the consortium.

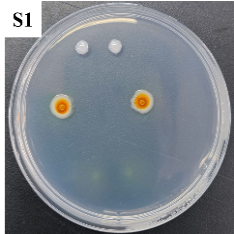

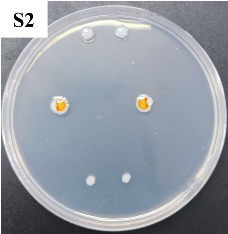

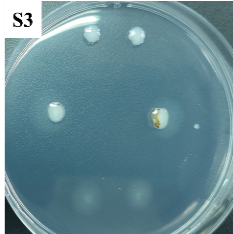

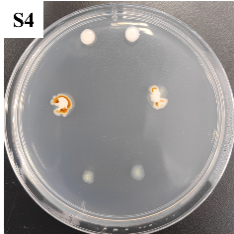

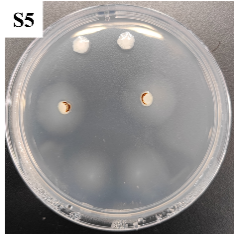

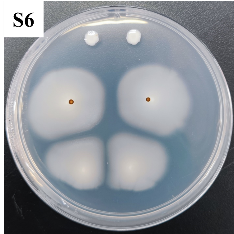

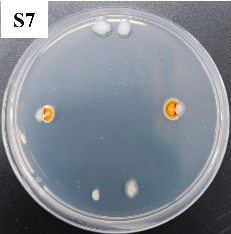

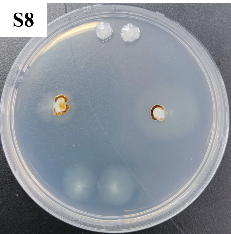

**Figure S6. Growth dynamics and Mn(II) oxidation in co-culture of *Arthrobacter* and** ***Tetrahymena thermophila* B2086.** (a) *Arthrobacter* colony forming unit (CFU) counts over time. A-mono denotes *Arthrobacter* monoculture; A-co denotes *Arthrobacter* and *T. thermophila* co-cultivation. (b) *T. thermophila* cell counts over time. T-mono denotes *T. thermophila* monoculture; T-co denotes *T. thermophila* B2086 and *Arthrobacter* co-cultivation. (c) Mn(II) oxidation under different culture conditions. A+T denotes Mn(II) oxidation by *Arthrobacter* and *T. thermophila* co-culture; A-mono denotes Mn(II) oxidation by *Arthrobacter* monoculture; T-mono denotes Mn(II) oxidation by *T. thermophila* monoculture.

**Experimental procedure:** *Tetrahymena thermophila* B2086 was kindly provided by Prof. Guangbo Qu (Research Center for Eco-Environmental Science, Chinese Academy of Sciences, Beijing, China). Cultures were maintained at 28°C in pathogen-free medium containing 2% protease peptone (Becton, Dickinson and Company, USA), 0.1% yeast extract (OXOID, Thermo Fisher Scientific, USA), 0.2% glucose (Sigma, USA), and 0.003% ferric citrate (Sigma, USA). Cultures were gently shaken at 135 rpm during incubation, and 1% (v/v) penicillin-streptomycin solution (HyClone, GE Healthcare Life Sciences, USA) was used to prevent bacterial or fungal infection and contamination. For co-culture assays, *T*. *thermophila* and *Arthrobacter* were washed and resuspended in phosphate-buffered saline (PBS), then adjusted to 5 × 10^4^ cells/mL and 10^6^ CFU/mL in PYG medium, respectively. Co-incubation experiments were performed in 100-mL flasks containing 30 mL of co-culture for 48 hours. Samples were taken every 12 h to quantify bacterial and protist populations and to measure residual Mn(II) concentrations. *Arthrobacter* populations were quantified via CFU counts on PYG agar plates, while *T.* *thermophila* counts were determined using a hemocytometer under an inverted microscope. Data represent means of three biological replicates. Error bars denote standard deviation.

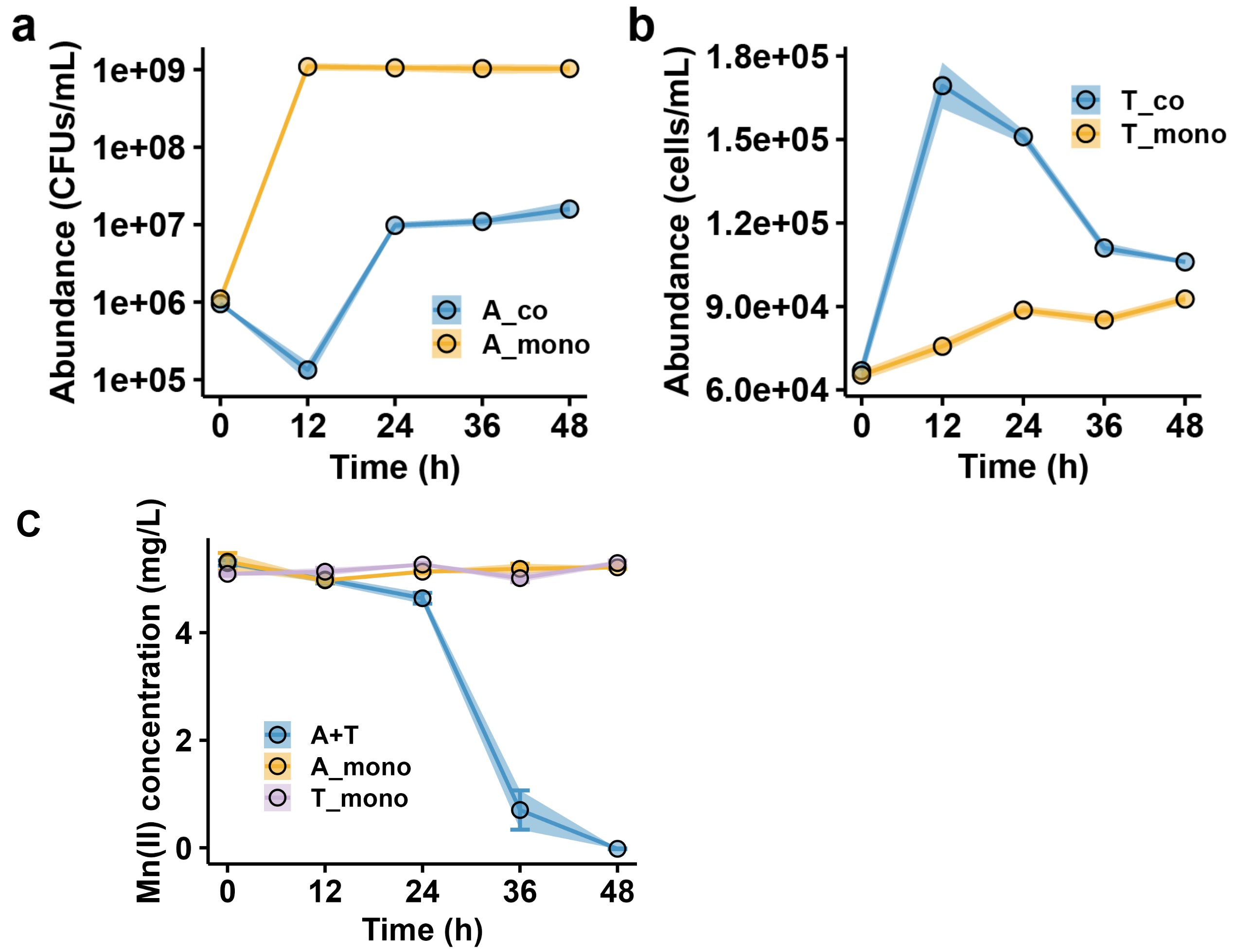

**Figure S7. Antibiotic-induced Mn(II) oxidation in *Arthrobacter*.** OTC: oxytetracycline; TC: tetramycin; OTC+A: simultaneous addition of oxytetracycline and *Arthrobacter*; TC+A: simultaneous addition of tetramycin and *Arthrobacter*.

**Experimental procedure:** Overnight cultures of *Arthrobacter* were incubated in 30 mL of PYG medium at 30°C and 170 rpm. These cultures (OD_600_ = 0.1, 10^8^ CFU/mL) were twice washed with PBS buffer and inoculated at 1% (v:v) into 2 mL of PYG medium containing oxytetracycline and tetracycline in 12-well plates. Mn oxide formation was visually monitored at 12-hour intervals. To identify the optimal concentration for Mn(II) oxidation induction, a gradient of antibiotic concentrations was tested: 10, 5, 2.5, 1.25, 0.625, 0.3125, 0.1563, 0.0781, and 0.0391 mg/L. Oxidation was most efficiently induced at 0.1563 mg/L for both antibiotics, with Mn(II) oxidation observed at 24 h. Higher or lower concentrations either delayed the response or produced no significant oxidation. This concentration (0.1563 mg/L) was used in subsequent assays, and photographs were taken after 48 hours of incubation.

**Table S1. Single bacterial strains capable of inducing Mn(II) oxidation in *Arthrobacter* under co-cultivation conditions.**

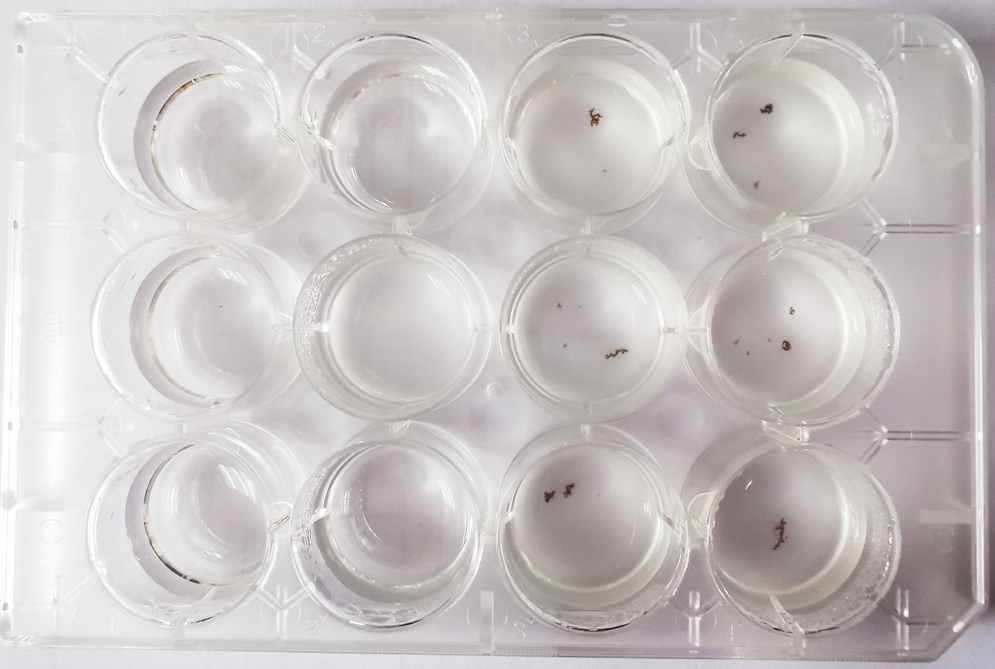

**TC**

**OTC+A**

**OTC**

**TC+A**

| Strain ID | Taxon | Accession no. of genome | Isolation source |
| --- | --- | --- | --- |
| H1 | *Lysobacter soli* | JARQWX000000000 | Grass soil |
| D1 | *Caulobacter segnis* | JARQWW000000000 | Forest soil |
| N1 | *Caulobacter endophyticus* | JARQWV000000000 | Mountain soil |
| S6 | *Sphingobium yanoikuyae* | JARQWY000000000 | Rhizosphere soil |
| X | *Stenotrophomonas maltophilia* | JARPOH000000000 | Riverside soil |
| X2 | *Dyella jiangningensis* | JARPOG000000000 | Riverside soil |
| X3 | *Sphingomonas* sp. | JARQXA000000000 | Grass soil |
| X6 | *Stenotrophomonas* sp. | JARQXB000000000 | Mountain soil |

**Table S2. External environmental stressors that trigger Mn(II) oxidation in *Arthrobacter***

| Stress type | Colony ID | Taxa in colony or chemical type | Accession no. of genome |
| --- | --- | --- | --- |
| Bacterial consortium | S1 | *Caulobacter rhizosphaerae*, *Flavobacterium amnicola* | Not submitted |
|  | S2 | *Pseudoduganella guangdongensis*, *Massilia eburnea*, *Duganella fentianensis* | Not submitted |
|  | S3 | *Stenotrophomonas nematodicola*, *Enterococcus avium* | Not submitted |
|  | S4 | *Stenotrophomonas maltophilia*, *Xanthomonas campestris* | Not submitted |
|  | S5 | *Stenotrophomonas rhizophila*, *Xanthomonas campestris* | Not submitted |
|  | S6 | *Massilia eburnea*, *Pseudoduganella guangdongensis*, *Phenylobacterium conjunctum* | Not submitted |
|  | S7 | *Stenotrophomonas maltophilia*, *Massilia agri* | Not submitted |
|  | S8 | *Stenotrophomonas maltophilia*, *Stenotrophomonas pavanii*, *Luteimonas cucumeris* | Not submitted |
| Eukaryote | / | *Tetrahymena thermophila* | / |
| Antibiotic | / | Tetramycin | / |
|  | / | oxytetracycline | / |
